# The borders of *cis*-regulatory DNA sequences harbor the divergent transcription factor binding motifs in the human genome

**DOI:** 10.1101/383182

**Authors:** Jia-Hsin Huang, Ryan Shun-Yuen Kwan, Zing Tsung-Yeh Tsai, Huai-Kuang Tsai

## Abstract

Changes in the *cis*-regulatory DNA sequences and transcription factor (TF) repertoires provide major sources that shape the gene regulatory evolution in eukaryotes. However, it is currently unclear how dynamic change of DNA sequences introduce various divergence level of TF binding motifs in the genome over evolutionary time. Here, we estimated the evolutionary divergence level of the TF binding motifs, and quantified their occurrences in the DNase I hypersensitive sites. Results from our *in silico* motif scan and empirical TF-ChIP (chromatin immunoprecipitation) demonstrate that the divergent motifs tend to be introduced at the borders of the *cis*-regulatory regions, that are likely accompanied with the expansion through evolutionary time. Accordingly, we propose that an expansion by incorporating divergent motifs within the *cis*-regulatory regions provides a rationale for the evolutionary divergence of regulatory circuits.

## Introduction

Transcription factors (TFs) are primary regulators of gene expression by interacting with DNA in a sequences-specific manner. The capability of a TF recognizing particular patterns of nucleotides *(i.e.* motif) via its DNA binding domains is defined as DNA-binding specificity [1]. Previous studies have reported that the DNA-binding specificities of the TF orthologs between human and Drosophila are mostly conserved [2]. Nonetheless, TFs do evolve divergent binding specificities in different species due to evolutionary variances, such as gene duplication and expansion of gene families [2–4]. Divergence in the TF binding specificities contributes significantly on differential gene regulation to shape the eukaryotic evolution[5–7].

In eukaryotic cells, multiple TFs cooperatively interact with genomic DNA to temporally and spatially regulate gene expression. Most eukaryotic chromatins are packed into nucleosomes and functional TF binding sites tend to be nucleosome depleted, whereby DNA is hypersensitive to cleavage by DNase I. DNase I hypersensitive sites (DHSs) have been studied extensively to be overlapped with most TF binding sites (TFBSs) in diverse organism. The major advance of ENCODE project has used DHSs to map active *cis*-regulatory elements in the human genome [8,9]. Integrative analyses using ENCODE data have identified hundreds of TF binding motifs [10,11] and extended the repertoire of TFs in the human genome [12]. And yet, there is high turnover in the *cis*-regulatory sequences [13], and over longer timescale, rapid and flexible TFBS gain and loss events occur between closely related species (14)–(16).

From a functional genomics perspective, the interplay between TF binding events and *cis*-regulatory regions is a pivotal step so that transcriptional regulation can be rewired through evolutionary time. General property of regulatory genomes lays on the broad presence of clustered TFBSs in the *cis*-regulatory regions [17,18]. The divergence of *cis*-regulatory sequences for harboring variable TFBSs but not alternations of TF binding motifs has been proposed as the major driving force to cause the phenotypic changes [19,20]. However, the manner in which the changes of DNA sequences in the *cis*-regulatory regions by harboring diversified TF binding motifs remains unclear. Since a given region of DNA sequences can harbor more than one TF binding motifs, the evovability within a *cis*-regulatory DNA sequences for various TF binding motifs has not been systematically studied.

In the present study, we have developed a measurement, motif prevalence index (MPI), for the divergent level of motifs among eukaryotes based on the discovery of general conservation of TF binding motifs among diverse organisms. The method integrates the phylogenetic relationship between TF orthologs among animals and a comprehensive collection of TF binding motifs from Cis-BP database [4], which provides a stringent inference for TF binding motifs among diverse organisms, to compute how prevalence of the human motifs across metazoan evolution. By averaging the MPI of all different motifs in the DNA region, we can study the evolution of DNA sequences for their preferences of TF DNA-binding motifs. Because the divergence of novel motifs could be defined as the presence in the later lineage of animals such as primate lineage, a given DNA sequences could be assessed weather more divergent motifs occurred. Our results showed the preference of the divergent motifs tend to locate in the borders of the open-chromatin regions. Furthermore, an integrative analysis using DHSs regions with TF chromatin immunoprecipitation (ChIP) sequencing from ENCODE project further confirmed our *in silico* results. Taken together, the discovery of introducing divergent motifs across evolutionary time would highlight the co-evolution between TF binding specificities and the functional effects of *cis*-regulatory variants on gene expression and therefore phenotypic evolution.

## Results and Discussion

### Motif Prevalence Index estimates the divergence level of motif sequences

We proposed a new measurement, Motif Prevalence Index (MPI), to estimate the evolutionary divergence level of TF DNA-binding preferences (motifs) in humans, according to the finding that the primary DNA binding specificities of TFs with similar DNA binding domain (DBD) sequences are generally conserved between distantly related species [2–4]. Based on phylogenetic distance and existence of a given motif *(i.e.* having homologous TFs with conserved amino acid sequences of DBD from the Cis-BP database) across metazoan species, MPI represented the evolutionary divergence level of the human motifs with a score from 0 (human-specific) to 1 (common in 74 metazoan species). Next, we selected the human motifs with experimental evidence from the JASPAR database [23]. Most of the human motifs (72.8% of the 364 motifs shown in Supplementary Table 1 and 2) were commonly presented across Metazoa and Bilateria taxa, whereas the divergent motifs (MPI < 0.1, 7.7%) in humans emerged approximately after the divergence of the vertebrata lineage (Figure 1). MPI was not biased by a couple of intrinsic motif properties, such as motif length and information content (no significant correlation as shown in Supplementary Figure S1). But the GC content was significantly lower in more divergent motifs (Supplementary Figure S1). Moreover, no significant correlation between MPI and the gene age of their corresponding TFs reflects the independence between the evolutionary history and the changes of binding specificity of TF repertoires.

**Figure 1.**
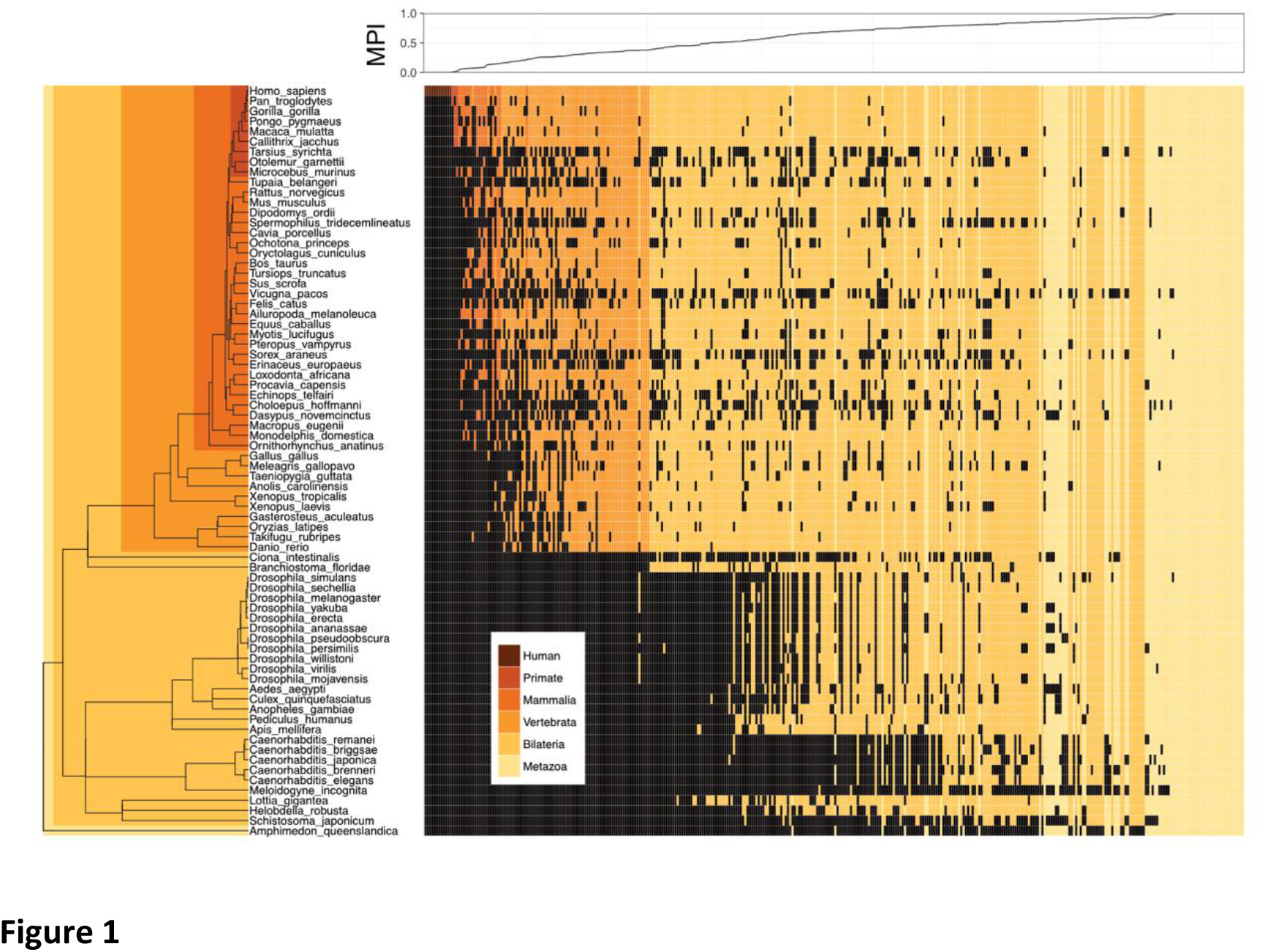
Motif prevalence index (MPI) of the TF binding specificities in humans. Phylogenetic relationship of 364 human TF binding specificities (motifs) from the JASPAR database and their MPI scores. Color codes denote the presence of motifs in different metazoan lineages. Black denotes the absence of motifs.

### Borders of DHS regions prefer divergent motifs

A theoretical study has suggested that the neighboring DNA sequences of the pre-existing TF binding sites (TFBSs) are preferred for the emergence of newly evolved binding sites [30]. Accordingly, we propose that the relatively common motifs are located around the middle of the open-chromatin regions whereas relatively divergent motifs are located at the border regions. To test this hypothesis, we conducted an *in silico* motif scan in 1-kb upstream to 500-bp downstream of transcription start site (TSS) of protein-coding genes and further filtered the DNase I hypersensitivity site (DHS) clusters in 125 cell types, which are highly corresponding to TFBSs [8]. We then investigated the open-chromatin regions as defined by the DHS peaks in the range of 150 – 400 bp in accordance with one to two nucleosome-free regions, which contain several TFBSs theoretically, and then computed the mean MPI of occurred motifs. Of note, to reduce the ambiguity of motif occurrences from similar motif pattern, we focused on 93 non-redundant JASPAR motifs that were clustered by Tomtom [24] with a threshold of p-value at < 0.05 and the MPI of selected motifs remained smoothly distributed (Supplementary Figure S2). In agreement with expectation, the spatial distribution of the mean MPI scores significantly decreased from center to border within the DHS regions (Figure 2A, Spearman’s correlation coefficient *rho* = –0.753, *p* < 2.2 × 10^−16^). Specifically, the mean MPI scores in the DHS-edge (i.e. decile regions of both DHS borders) were significantly lower than those in the DHS-center (i.e. quintile regions of DHS center) (Figure 2A, one-sided Wilcoxon rank-sum test, *p* = 4.76 × 10^−30^). In contrast, the closed chromatin regions in the promoters not only possessed higher mean MPI scores than open chromatin regions but also showed a negligible decline in the mean MPI scores (Figure 2A, Spearman’s correlation coefficient *rho* = –0.01). Since the divergent motifs with lower MPIs are the TFs that have evolved to recognize new DNA sequences across evolution, a question immediately arose is whether the DNA sequences in the DHS regions show distinct conservation levels.

**Figure 2.**
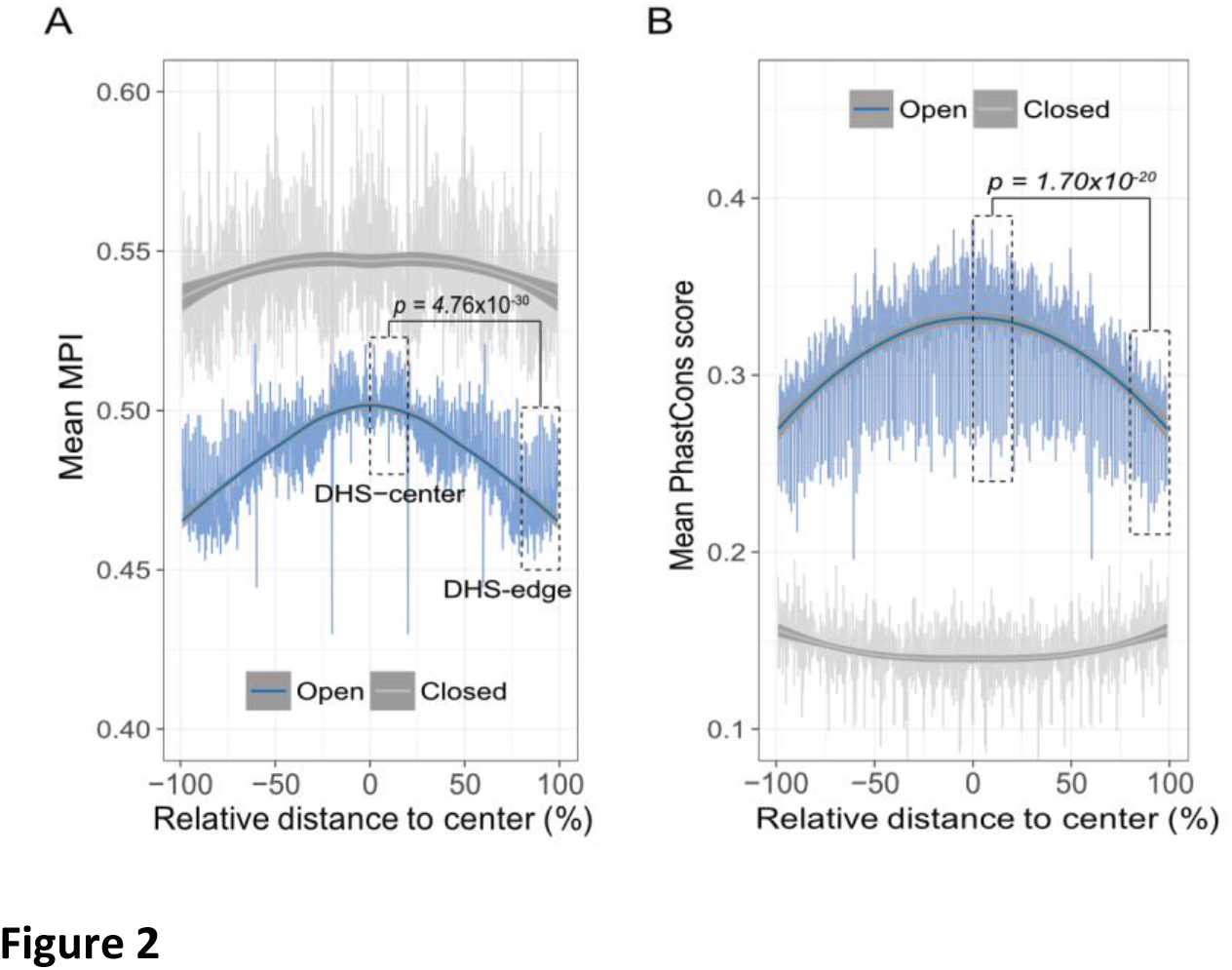
The borders of open chromatin regions for the emergence of the divergent TF binding motifs. (A) Distribution of mean MPIs at relative positions within the chromatin regions between 150 to 400 bp. Since DHS regions having different lengths of peaks, the mean MPI distribution was calculated in 0.1%relative distance sliding windows for DHSs. The relative distance was defined as the normalized distance from the center of the fragments, ranging from 0% at the center to 100% at the edge of a given DHS peak. The mean MPI scores mirror each other around the center of the DHS regions. DHS-center denotes the quintile regions of DHS center; DHS-edge denotes the decile regions of both DHS borders. A p-value between DHS-center and DHS-edge was obtained by one-sided Wilcoxon rank-sum test. Open denotes the DHS regions and closed denotes the promoter regions without overlapping with DHSs. (B) Distribution of the mean PhastCons conservation scores at relative positions within the open and closed chromatin regions between 150 to 400 bp. A p-value between DHS-center and DHS-edge was obtained by one-sided Wilcoxon rank-sum test.

Thus, we sought to determine whether the decreasing pattern of mean MPIs along position was systematically paralleled by patterns of evolutionary conservation in open-chromatin regions. We used the PhastCons score [31] to calculate the levels of evolutionary conservation of DNA sequences from alignments of 99 vertebrate genomes [32]. As expected, the open chromatin regions (DHSs) possessed higher conservation levels than the closed chromatin regions (Figure 2B). However, the PhastCons scores were significantly higher in the DHS-center than in the DHS-edge (Figure 2B, one-sided Wilcoxon rank-sum test, *p* = 1.70 × 10^−20^). That is, less evolutionary constraint at the DHS borders may reflect the rapid TFBS turnover, introducing the divergent motifs readily.

Theoretically, regulatory complexity such as the number of TF regulating a gene increases continuously over the evolutionary time [33]. We thus examined whether the differences in the mean MPI scores between DHS-center and DHS-edge remained in different ages of genes. The results showed that the significant differences on the mean MPI scores between DHS-center and DHS-edge were consistent in the promoters of all ages of genes (Figure 3A). Nevertheless, we noticed that the numbers of longer DHSs increased in the older group of genes (Supplementary Figure S3). We then performed a further analysis (Figure 3B) according to different lengths of DHSs and found that the differences on the mean MPI scores between DHS-center and DHS-edge increased in the longer DHSs (> 200 bp). Intrigued by these results, we assessed the fold enrichment of occurrences between divergent (MPI < 0.1) and common motifs (MPI >= 0.9) across gene ages and DHS lengths. The results showed that the divergent motifs were not enriched in the short DHS (150-199 bp) regions but enriched at the border regions of longer DHSs (Figure 3C). Similar robust results were found when applying different cut-offs for specific (MPI < 0.2) and common motifs (MPI >=8) (Supplementary Figure S4). Therefore, one feasible interpretation for our observations is that the introduction of divergent motifs is likely to accompany with the elongation of the *cis*-regulatory DNA regions, specifically on the boundaries. With increased number of longer DHSs in the promoter of the older gene, such expansion of *cis*-regulatory regions with introduction of divergent motifs could contribute to the regulatory complexity of genes across evolutionary time.

**Figure 3.**
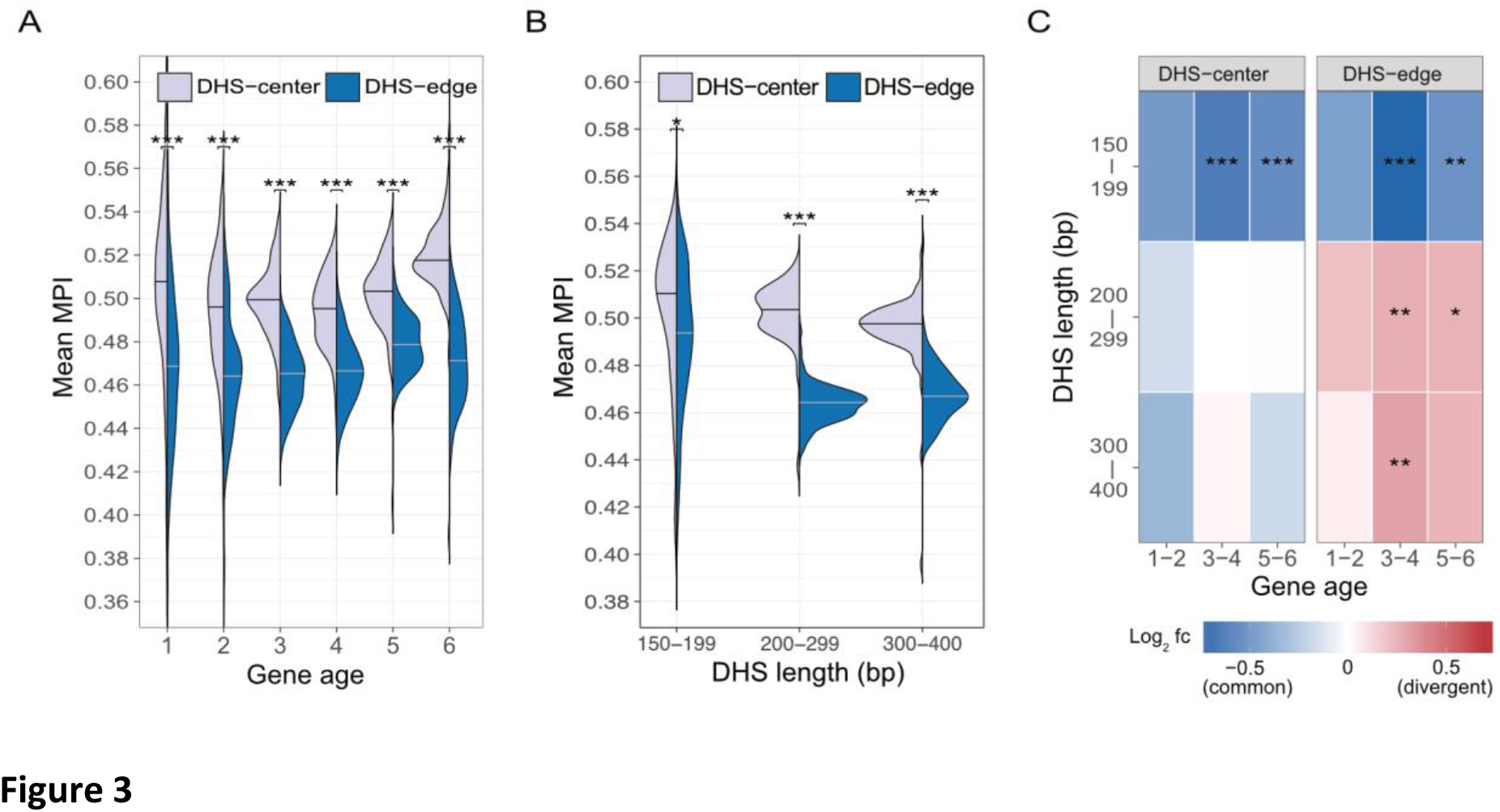
Motif enrichment in the promoter regions of protein-coding genes across different gene age and DHS lengths. (A) Comparison of mean MPIs between DHS-center (purple, left violin plot) and DHS-edge (blue, right violin plot) in six age categories of genes. The age of human genes arose at different evolutionary time were identified by combining homolog clustering with phylogeny inference according to Yin*et al.* 2016 [47]. Accordingly, category 1 to 6 denoted the Primate origin (youngest genes), Mammalia, Vertabrata, Metazoan, Eukaryota; and Cellular organism (oldest genes), respectively. (B) Changes of mean MPI between DHS-center and DHS-edge by the DHS length. The DHSs lengths were confined to specific widths (150–199, 200–299, 300–400 bp). Significant values in (A) and (B) were obtained by Wilcoxon rank-sum test after Bonferroni correction for multiple tests. (C) Enrichment for motif occurrences. Color code in the cells indicates the fold-changes *(Log2* fc) of occurrences of the divergent motifs divided by common motifs. Divergent motifs were MPI < 0.1; common motifs were MPI >= 0.9. Fisher’s exact test was applied to examine whether the proportion was significantly different (2 × 2 contingency table where rows correspond to occurrences inside/outside of the part, and columns represent TF groups). Significant values were obtained after Bonferroni correction for multiple tests. Note that *: *p-value* < 10^−2^, **:*p-value* < 10^−3^, ***: *p-value* < 10^−4^.

### TF ChlP-seq reveals similar distribution of MPI within DHS regions

To further validate our discovery of motif distribution within the *cis*-regulatory DNA regions independently from the motif scan approach, we conducted our analysis by overlapping DHSs with *in vivo* chromatin immunoprecipitation followed by DNA sequencing (ChIP-seq) of 243 TFs (Supplementary Table S1) downloaded from the ENCODE project [17], and then recalculated the mean MPI scores by the corresponding MPIs of TFs. Remarkably, the empirical TF-ChIP results displayed a significant decline of the mean MPI scores from center to border within the DHS regions in a genome-wide scale (Fig. 4A, Spearman’s correlation coefficient *rho* = −0.940, *p* < 2.2 × 10^−16^). This result was highly consistent with the *in silico* motif scanned results (Figure 2A). Additionally, the mean MPI between DHS-center and DHS-edge were significantly different among different *cis*-regulatory regions such as promoters of other genes (protein-coding genes, non-coding genes, and pseudogenes) and enhancers, which were obtained from FANTOM5 [34] (Supplementary Table S3). For different lengths of DHSs, the TF-ChIP results also confirmed the significant differences on the mean MPI scores between DHS-center and DHS-edge (Figure 4B).

Besides, we noticed that the motifs corresponding to those pioneer TFs, which were reported for the chromatin-remodeling activity [35], had significantly higher MPIs than others (Figure 4C, one-sided Wilcoxon rank-sum test, *p* = 2.52 × 10^−3^). Since the pioneer TFs have been known to disrupt chromatin structure to create a nucleosome-free DNA region and thus open the nearby regions allowing other TFs to access DNA [36,37], the binding of pioneer TFs with higher MPI provides a likely rational for the spatial distribution of mean MPI in the DHS regions. Therefore, our observations from the results of *in silico* motif scan and empirical TF-ChIP unveiled a differential preference within the *cis*-regulatory DNA regions, where the regions evolved motifs bound by TFs with different divergent levels of binding specificities.

### TFs with divergent motifs tend to express ubiquitously among human tissues

Based on the expression profiles in 32 human tissues from Human Protein Atlas (HPA) [29], TFs could be divided into one group showing a ubiquitous expression in most tissues and another group showing a significantly elevated expression in at least one of the human tissues. Remarkably, the majority of TFs possessing more divergent motifs are ubiquitously expressed in the human tissues, whereas the fraction of TFs possessing common motifs displays an elevated expression pattern in specific human tissues increased with MPI (Figure 4D for the TFs with ChIP-seq, Supplementary Figure S5 for all other TFs from HPA). Of note, a recent study has reported that the duplicate genes tend to diverge in their expression profiles among tissues across evolution [38]. According to our observations, a common motif usually corresponds to several members of the TF paralogs (Supplementary Table S1). This increased fraction of TFs showing tissue-elevated expression most likely accounted for the expansion of gene paralogs. Thereafter, we computed the fold enrichment of TF-ChIP peaks within the DHS regions by comparing the ubiquitously expressed TFs with divergent motifs (MPI < 0.1) to the TFs with common motifs (MPI >= 0.9) that showed either ubiquitous expression or tissue-elevated expression. We found that the ubiquitously expressed TFs with divergent motifs were significantly enriched at the DHS-edge and had a higher proportion than with the common motifs (Figure 4E, Supplementary Figure S6 for the enrichment analyses). In contrast, the tissue-elevated TFs with common motifs had the highest proportion and significantly enriched at the DHS-center (Figure 4E, Supplementary Figure S6). Taken together, these results confirm our proposed hypothesis and imply another level of the dynamics of transcriptional regulation on the interplay of DNA motifs and distinct expression patterns of TFs.

**Figure 4.**
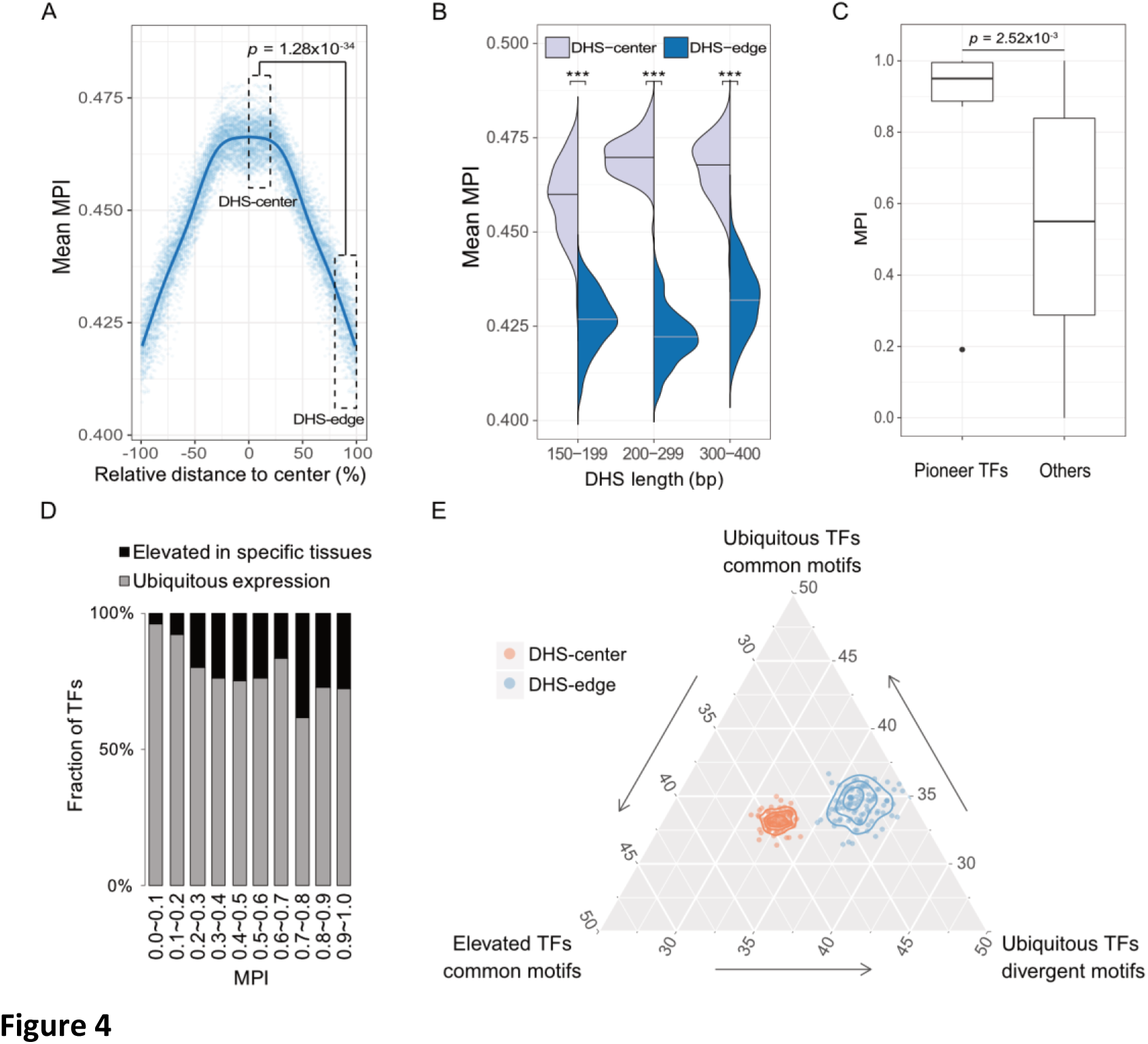
The differential preference of the *cis*-regulatory regions harboring the TF binding motifs in the human genome. (A) Distribution of mean MPIs at relative positions of DHSs based on the overlapped ChIP-seq peaks of 243 TFs with the genome-wide DHS regions, whose lengths were between 150 to 400 bp. The mean MPI scores mirror each other around the center of the DHS regions. A p-value between DHS-center and DHS-edge was obtained by one-sided Wilcoxon rank-sum test. (B) Changes of mean MPI between DHS-center and DHS-edge by different DHS lengths. Significance were obtained by one-sided Wilcoxon rank-sum test and followed by Bonferroni correction, *: *p-value* < 10^−2^, **: *p-value* < 10^−3^, ***: *p-value* < 10^−4^. (C) Differences of MPIs between motifs corresponding to pioneer TFs and other motifs and a p-value was obtained by one-sided Wilcoxon rank-sum test. (D) The fraction of the TFs with ChIP-seq classified according to the tissue expression pattern (Uhlén et al., 2015) for each of corresponding MPI ranges. Grey denotes the ubiquitous expression in most humantissues and black denotes the elevated expression in the specific tissues. (E) The ternary proportion distributions for the TF-ChIP occurrences in the DHS regions. The proportion values were determined by the fraction of each group of TF-ChIP occurrences in the given DHS regions. The ubiquitous TFs with divergent motifs (MPI < 0.1, ubiquitous expression) and the TFs with common motifs (MPI >= 0.9, ubiquitous expression or tissue-elevated expression) were grouped as in Fig. 4D. Center denotes the quintile regions of DHS center; edge denotes the decile regions of both DHS borders.

The DNA sequences in the *cis*-regulatory regions are dynamic for harboring different TFs. Through the changes in the mean MPI scores, which corresponded to the DNA sequences using different divergence levels of TF binding specificities, the borders of *cis*-regulatory regions are more preferred for introducing divergent motifs than the center regions (Figure 2A, 4A). Our results are in line with the theoretical studies, which show the sequences adjacent to pre-existing TFBSs readily evolve for the emergence of new TFBSs [30,39]. As common motifs with high MPI are prevalent among metazoan species, the center region of *cis*-regulatory regions are most likely to be ancestral binding sites and constrained over evolutionary time as shown by higher PhastCons scores (Figure 2B).

Finally, we proposed a model for the expansion of TFBSs with conserved motifs by introducing divergent motifs to adjacent sites in the *Cis*-regulatory regions (Figure 5). *cis*-regulatory evolution, such as changes in the TFBSs over evolutionary time scale, is an important source for the diversity of morphological traits through gradual modification of transcription circuits (40)–(42). Since TFs often bind to adjacent sites of regulatory regions cooperatively [43,44], the regulatory circuits, through coordinating alternative TFs, could diversify as that motifs on the TFBS-clustered border regions could be replaced for expansion of new motifs. Furthermore, the center region of *cis*-regulatory regions is even highly intertwined with many TF paralogs that are particularly with a tissue-elevated expression. Since rewiring of regulatory networks is crucial for divergent expression patterns in evolution [45,46], we suspect that an expansion mechanism by incorporating more divergent motifs at the borders of *cis*-regulatory regions serves as common evolutionary intermediates in rewiring regulatory networks.

**Figure 5.**
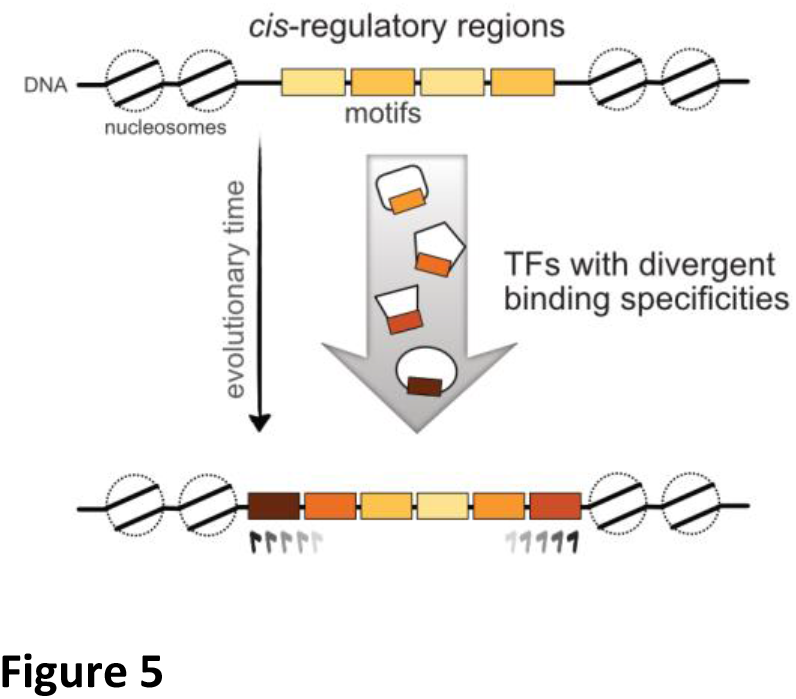
The proposed model for the dynamics of TF binding motifs in the *cis-* regulatory regions. The borders of the *cis*-regulatory regions are preferred for harboring the divergent motifs across evolutionary time.

## Materials and Methods

### Motif Prevalence Index

The primary TF binding motifs of human and 73 metazoan species were obtained from the Cis-BP database [4]. Given a motif *x, n* species *S_1‥n_* possessing its corresponding TF families can be revealed based on annotations in the Cis-BP database. We then constructed a phylogenetic tree *T_S_* of *n* species *S_l…n_* by applying neighbor-joining method [21] to evolutionary distances from the TimeTree database [22] between each pair of species among *S_l…n_*. Given *B(T)* is the total length of branches in a phylogenetic tree T, Motif Prevalence Index (MPI) was defined as the ratio of the length sum of the branches of *TS* to the length sum of the branches of the upper-limit tree of 74 metazoan species, *B(T_S_)/B(T_74 metazoan_)*, which is a score between 0 and 1. To obtain a non-redundant and reliable TF set for the matrix-scan analysis, we selected 364 motifs which were well curated TF models from JASPAR 2018 database [23]. We further applied Tomtom [24] to group them into 93 clusters of non-redundant motifs with a threshold of *p*-value < 0.05, and then the motifs possessing the highest MPI of each cluster were retained.

### Identification of TF binding motifs in open and closed chromatin regions

The human genome sequence and gene annotation were obtained from Ensembl (GRCh37, release 75) [25]. We identified the occurrences of TF binding motif in promoter regions (−1k to +500 bp from TSS) for each of the 93 motifs by scanning its position probability matrix using Matrix-scan of RSAT tool box [26] with a threshold of false discovery rate < 10^−4^. DNase I hypersensitive (DHS) cluster data were downloaded from the UCSC genome browser [27] for 125 cell types determined by ENCODE project [8]. DHS peaks were defined as open chromatin regions and the chromatin region without overlapped DHS peaks were defined as closed chromatin regions.

### Transcription factor ChIP-seq datasets

The ChIP-Seq peaks of 243 TFs (Supplementary table S1) in numerous cell lines were downloaded from ENCODE Consortium [28] based on genome hg19 assembly. For each TF, the tracks of the same cell lines were combined by retaining the overlapping base pairs with at least half of the tracks. Since averaged length of ChIP-seq peaks were longer (~300 bp) than that of the TF binding motif, we applied TF binding sites as 25 bp before and after the summit of ChIP-seq peaks, respectively.

### The expression pattern of TFs

The expression profile of the human TF were according to the Human Protein Atlas (HPA) [29]. Since HPA has defined five categories of all human expressed genes, we grouped the expression of TF genes in relatively general terms as ubiquitous expression and tissue-elevated expression in the present study. The categories of expressed in all tissues and mixed from HPA were denoted as ubiquitous expression. The categories of tissue enhanced, group enriched, and tissue enriched from HPA were denoted as tissue-elevated expression.

## Code Availability

The computer codes that support the findings of this study are available in Git-Hub with the identifier doi:10.5281/zenodo.1208608.

## Acknowledgements

This work was supported by the Institute of Information Science, Academia Sinica and the Ministry of Science and Technology (MOST 106-2811-E-001-005 to J.-H.H. and MOST 105-2221-E-001-029-MY3 to H.-K.T.).

## Author contributions

J.-H.H. and H.-K.T. conceived the idea, designed the study and wrote the manuscript. S.-Y.K. and Z.T.-Y.T. developed the computational algorithms and performed the bioinformatics analysis. Z.T.-Y.T. provided guidance in data analysis and interpretation of the results. All authors contributed to amending the manuscript and have read the submitted version.

